# Subtle Visual Latency Can Profoundly Impair Implicit Sensorimotor Learning

**DOI:** 10.1101/2024.03.14.585093

**Authors:** Alkis M. Hadjiosif, George Abraham, Tanvi Ranjan, Maurice A. Smith

## Abstract

Short sub-100ms visual feedback latencies are common in many types of human-computer interactions yet are known to markedly reduce performance in a wide variety of motor tasks from simple pointing to operating surgical robotics. These latencies are also present in the computer-based experiments used to study the sensorimotor learning that underlies the acquisition of motor performance. Inspired by neurophysiological findings showing that cerebellar LTD and cortical LTP would both be disrupted by sub-100ms latencies, we hypothesized that implicit sensorimotor learning may be particularly sensitive to these short latencies. Remarkably, we find that improving latency by just 60ms, from 85 to 25ms in latency-optimized experiments, increases implicit learning by 50% and proportionally decreases explicit learning, resulting in a dramatic reorganization of sensorimotor memory. We go on to show that implicit sensorimotor learning is considerably more sensitive to latencies in the sub-100ms range than at higher latencies, in line with the latency-specific neural plasticity that has been observed. This suggests a clear benefit for latency reduction in computer-based training that involves implicit sensorimotor learning and that across-study differences in implicit motor learning might often be explained by disparities in feedback latency.

## Introduction

Visual feedback latencies are an inherent component of human-computer interactions that rely on continuous task feedback, from everyday computer mouse use^1^ to sophisticated virtual and augmented reality systems used for skill training in tasks like the operation of surgical robotic systems^2,3^, rehabilitation^4^, and flight simulators^5^. While these latencies due to delayed system response times are commonly short (<100ms) – a level at which they are often not perceived^6^– research has shown that even sub-100ms feedback latencies can markedly reduce motor performance in tasks such as reaching, tracking, steering, and collaborative control^7–12^.

These latencies are common in experiments that measure sensorimotor learning. Reliable reports range from a relatively low 36 ms^13^ to 60-80ms^14–16^ to 145ms^17^. However, latencies are unfortunately seldom measured, and values near or above 100ms are likely not uncommon based on personal communications with colleagues. Latency values above 100ms are not surprising given that many experimental setups use video projectors for visual feedback, and older projectors commonly inject display latencies in the order of 100ms^18^, that would add to latencies from sensor input, computer processing, and graphics output especially when common double-buffering schemes are used for graphics. The existence of these latencies raises the questions of whether experimental setups can often impair the very sensorimotor learning processes they try to measure, and whether between-setup differences in latency complicate the comparison of findings across different labs.

While the effect of visual feedback latency on sensorimotor learning has been the subject of multiple studies^13–15,17,19–31^, only a fraction^12–15,17,19^ reliably measured the baseline latency upon which additional experimentally-imposed delays were administered. Therefore, remarkably, both the actual latency of the experimental conditions and of the reference conditions to which they were compared were unknown in most studies. This is especially problematic for studies examining short sub-100ms delays^20,22,23^, but might not be a critical concern for studies that tested the effects of long experimentally-imposed delays ≥ 1000ms^27–31^ or even those with delays ≥ 200ms^21,24–26^, which are likely, in retrospect, to be well above the baseline latency, although we cannot be certain. Two long-delay studies dissected learning into implicit and explicit contributions and consistently found both decreased implicit and increased explicit learning following added delays of 200ms or greater^13,15^. But with no data at shorter delays, we cannot know whether these findings might apply to the sub-100ms latency range common in human-computer interactions including experiments for studying sensorimotor learning. This is especially the case because the few studies that examined small delays that might correspond to sub-100ms latencies (1) reported conflicting results, (2) failed, in all but one case, to measure actual latencies so that the studied latencies were not known, and (3) did not dissect learning into implicit and explicit contributions^19,20,22,23^. Two, including the only one to measure latencies^19^, reported that sensorimotor learning was not affected by added delays of up to 60ms^19,20^ whereas the other two, one of which was based only on data from a single animal, reported reduced learning for 50ms of added delay without measuring the actual latency^22,23^. It is therefore unclear how sub-100ms latencies, typical of human-computer interactions and motor learning experiments, might affect sensorimotor learning. If these small latencies can have large effects on sensorimotor learning, then experiment-to-experiment variability in latency within the sub-100ms range or just above it might explain the wide discordance in the amount of implicit or explicit motor adaptation observed. For example, even for studies using the same error-clamp paradigm to isolate implicit adaptation, the capacity for implicit visuomotor adaptation has been reported to be as low as 12° and as high as 25°, a greater than 2-fold difference^32–34^.

Moreover, evidence from neurophysiology suggests that latencies as small as 20ms can disrupt the neural plasticity that may underlie implicit sensorimotor learning. Both the spike timing-dependent plasticity (STDP) that mediates LTP in cortical neurons and the neural plasticity that mediates LTD in cerebellar Purkinje cells are governed by precisely-timed coincident input, with plasticity windows on the order of 20ms^35–40^. This leads to two possibilities regarding the effect of short latencies upon implicit sensorimotor learning. If sensorimotor learning relies on neurons tuned to a specific physiological latency associated with sensory input, then it should be exquisitely sensitive to short latencies. Alternatively, if sensorimotor learning could instead rely on one of several subpopulations of neurons tuned to a broad range of different preferred latencies, then it could be robust against changes in latencies provided they are within the distribution of these preferred latencies – which, for cerebellar Purkinje cells, is up to 150ms wide^40^. Previous work cannot distinguish between these two possibilities, as results are only consistent for delays ≥200ms^13–15,21,24–31^ corresponding to latencies well above 200ms, which lie beyond both the narrow tuning of individual cells and the width of the distribution of preferred latencies, thus predicting impaired learning in both cases.

To disambiguate between these two possibilities, we examined visuomotor learning with a short, 85ms visual feedback latency, comparing it against the 25ms optimized latency of our setup and a larger 300ms latency, as a long-latency reference. Using an aim report paradigm^15,41^, we dissected the learning measured on every training trial into implicit and explicit components, and found that implicit learning increased dramatically when latency improved by as little as 60ms, from 85ms to 25ms, whereas, explicit learning, by contrast, decreased, dramatically altering the balance between implicit and explicit adaptation. Remarkably, the sensitivity of implicit learning to these sub-100ms latencies was about 5-10 times greater than for higher latencies. These findings support the idea that implicit sensorimotor adaptation relies on neuronal populations narrowly tuned around a specific latency. The exquisite sensitivity of sensorimotor learning to sub-100ms latencies that we uncover may explain across-study differences in the amount of implicit adaptation that can be attained.

## Results

We examined the effect of short sub-100ms visual feedback latencies on sensorimotor learning by measuring the learning curves for the both implicit and explicit components of the adaptive response following exposure to a 30-degree visuomotor rotation (VMR) at different visual feedback latencies. Secondarily, we examined subsequent relearning following a block of no-feedback movements and measured the generalization of adaptive responses across different movement directions to provide additional insight into implicit and explicit learning. We studied three latency conditions: 25ms, 85ms, and 300ms. 25ms, measured with high-speed video, was the lowest visual feedback latency we could achieve following optimization of the experimental setup (see Methods) and thus served as our low-latency reference. To this reference, we added 60ms (a short additional latency) and 275ms (a longer additional latency similar to values tested in previous work^13,14,21,24,25^, used as a long-latency reference) in software to implement the 85ms and 300ms latency conditions, respectively, in order to investigate how sensorimotor learning is altered by sub-100ms vs longer latencies. In brief, participants (n=42) performed point-to-point reaching movements on a digitizing tablet, while receiving visual feedback in the form of an onscreen cursor that continuously tracked hand motion. The cursor was displayed on a computer monitor that was updated at 120Hz and positioned horizontally above the tablet (Fig. 1a). After a short task familiarization period where visual feedback was presented at a 25ms latency, the latency was set with equal probability to 25ms, 85ms, or 300ms for a 100-trial baseline period and subsequent 120-trial VMR training period, where participants moved in a single target direction while a fixed clockwise (CW) or counterclockwise (CCW) 30deg VMR (counterbalanced – see Methods) was applied. They then performed a generalization block of 114 trials in 19 different directions where visual feedback was withheld, followed by 60 retraining trials where visual feedback was provided with the same latency and the same VMR as in the 120-trial training period. This retraining block was followed by a second 114-trial no-feedback generalization block, and finally by a zero-rotation washout block of 50 trials where the latency was maintained, but the VMR was removed, to complete the superset. Participants then performed another superset of baseline, training, generalization, and washout blocks, but with a different visual feedback latency, a different target direction, and an oppositely signed VMR during training. The combination of the VMR inversion and the ±120° target direction change successfully minimized carryover effects from the previous superset. In particular, the difference between the baseline movement biases in the pre-training period at the beginning of supersets preceded by CW vs CCW VMR training was < 0.2° for the overall movement direction and also for implicit and explicit contributions to it, and not significantly different from zero (t<0.4 and p>0.7 in all 3 cases). Moreover, visual latency, target directions, VMR directions, and test order were independently balanced across participants within each experiment, so that any systematic effect of target direction, VMR direction or test order would be independent of the latency condition (see Methods). Experiments 1a (n=24) and 1b (n=18) were identical except that the target directions differed and participants were studied at two vs three latencies, respectively, i.e. each performed two vs three supersets of baseline / training / washout.

**Fig. 1:**
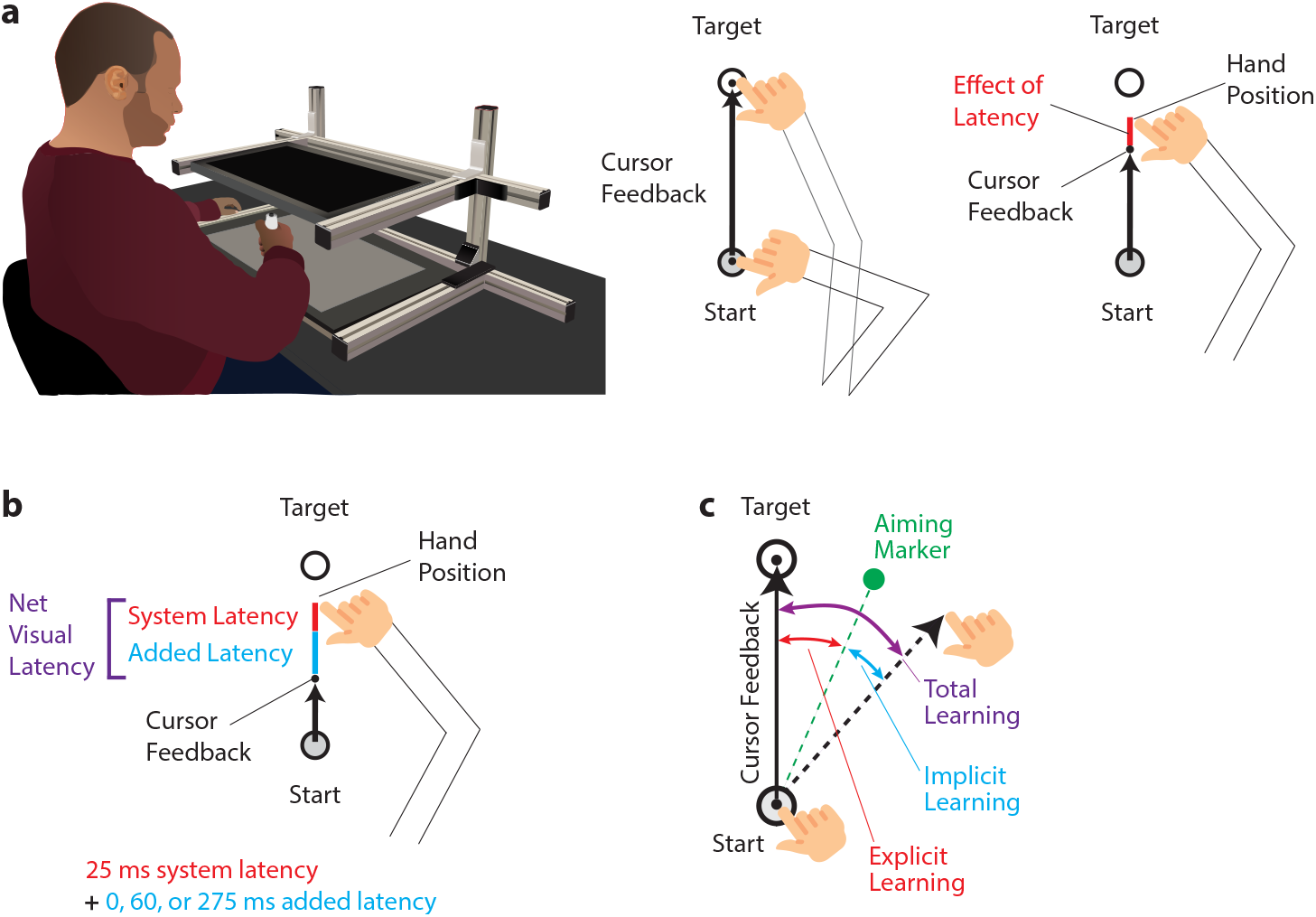
Visual latency in a reaching task. **(a)** -to-point reaching movements on a digitizing tablet while a cursor provided visual feedback on a horizontal screen positioned above the hand. Any latency in the visual display will make the cursor lag behind true hand motion. **(b)**The overall visual latency in our experiments consisted of a base system latency that was optimized down to a 25ms value that combined with experimentally-imposed delays of 0, 60 or 275ms to yield latencies of 25, 85 or 300ms. **(c)** Visuomotor rotation (VMR) with aiming paradigm. Users indicated their aim strategy before each movement by positioning an on-screen marker (green), allowing us to dissect total learning into explicit strategy vs implicit components (the angle between the aiming marker and target in red vs the angle between the hand motion and aiming marker in blue).

### Implicit sensorimotor adaptation increases and explicit strategy decreases when latency is reduced

Inspection of the learning curves for overall adaptation (shades of purple in Fig. 2a) reveals them to be remarkably similar in both shape and amplitude across all three tested latency conditions. **Correspondingly, when we quantified** overall sensorimotor adaptation by computing its asymptotic **level in the training block (operationally defined as the average adaptation over the last 20 trials of** the 120-trial training period, excluding trials following rest breaks, **see the grey “Late learning” regions** indicated in Fig. 2a-c), we found essentially identical overall adaptation levels for all three latency conditions (25ms vs. 85ms: 29.5±0.4° vs. 28.8±0.4° [mean±SEM], t(66)=1.2, p=0.11; 25ms vs. 300ms: 29.5±0.4° vs. 29.1±0.5°, t(66)=0.6, p=0.26, see Fig. 2d). However, decomposing adaptation into implicit and explicit components using aim reports^41,42^ (see Methods) revealed large opposing effects for implicit vs. explicit sensorimotor adaptation. Implicit adaptation was weakest in the 300ms condition but grew stronger as the latency of visual feedback decreased to 85ms and then 25ms (shades of blue in Fig. 2b). In contrast, explicit adaptation was strongest in the 300ms condition but grew weaker as the latency of visual feedback decreased (shades of red in Fig. 2c). Implicit adaptation was a remarkable 50% higher when latency was 25ms compared to 85ms (20.4±1.3° vs. 13.8±1.4°, respectively, t(66)=3.4, p=0.00049) and twice as high compared to the 300ms condition (20.4±1.3° vs. 10.9±1.6°, t(65)=4.7, p=6.6×10^-6^). In contrast, explicit adaptation was 40% lower when latency was 25ms compared to 85ms (9.1±1.3° vs. 15.0±1.5°, respectively, t(66)=3.0 p=0.0022) and 50% lower compared to the 300ms condition (9.1±1.3° vs. 18.1±1.3°, t(65)=4.9, p=3.9×10^-6^).

**Fig. 2:**
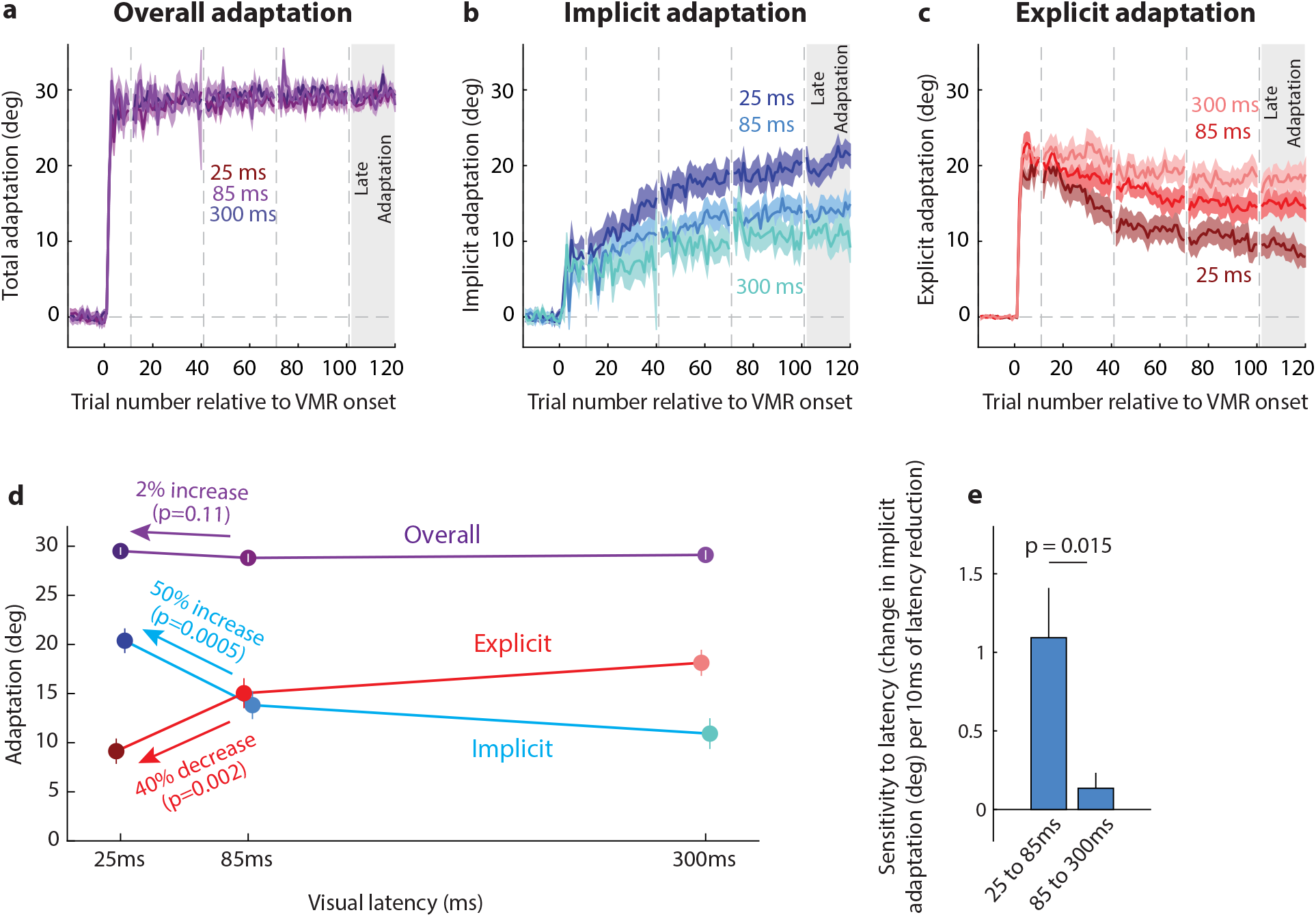
Small reductions in latency improve implicit and decrease explicit learning. **(a-c)** curves for **(a)** overall, **(b)** implicit, and **(c)** explicit adaptation for the three latency conditions studied shows increasing implicit and decreasing explicit learning as latency is reduced, with overall learning largely unaffected by latency. The gray rectangle indicates the late learning period analyzed in panel **d**. Vertical dashed lines indicate trials following 60s breaks. **(d)** overall learning is largely unaffected by latency, implicit learning is increased by 50% when latency is reduced from 85ms to 25ms and doubled when latency is reduced from 300ms to 25ms. In contrast, explicit learning is reduced by 40% when latency is reduced from 85ms to 25ms and by 50% from 300ms to 25ms. **(e)** Sensitivity of implicit learning to changes in latency. This sensitivity, the rate at which implicit learning increases per unit of latency decrease, is nominally 5-fold higher over the sub-100ms latency interval between 25 and 85ms compared to the interval between 85 and 300ms. Shading and error bars indicate ±SEM; note that error bars for overall learning in panel (d) are shown in white for visibility, as they would be occluded by the circle symbols otherwise.

A subsequent 60-trial retraining block within each superset, with a target direction, VMR, and latency that were all identical to the 120-trial training block, was separated from that training block by 114 no-feedback trials in different target directions (these trials were used to measure directional generalization - see Methods and the generalization analysis section later in the Results). This retraining block provides information about the effects of extended learning, as 50% more training was added to the initial training block. Analysis of sensorimotor learning in this block mirrors the 20 trials of the block to assess the asymptotic adaptation levels for overall, implicit, and explicit adaptation. We again observed little difference in overall adaptation between latency conditions (Fig. 3a,d), but found pronounced increases in implicit adaptation (Fig. 3b,d) and decreases in explicit adaptation (Fig. 3c,d) as latency was reduced. Implicit adaptation was in this case 60% higher when latency was 25ms compared to 85ms (19.9±1.3° vs. 12.4±1.7°, respectively, t(66)=3.5, p=0.00036) and again nearly double compared to the 300ms condition (19.9±1.3° vs. 11.1±1.7°, t(65)=4.1, p=5.6×10^-5^). In contrast, explicit adaptation was again 40% lower when latency was 25ms compared to 85ms (9.8±1.4° vs. 16.6±1.7°, respectively, t(66)=3.1, p= 0.0015) and 50% lower compared to the 300ms condition (9.8±1.4° vs. 18.5±1.4°, t(65)=4.3, p=2.9×10^-5^). Together these findings provide evidence for a dramatic increase in implicit sensorimotor learning alongside complementary decreases in explicit strategy arising from a small 60ms reduction in the latency of visual feedback in the sub-100ms range.

**Fig. 3:**
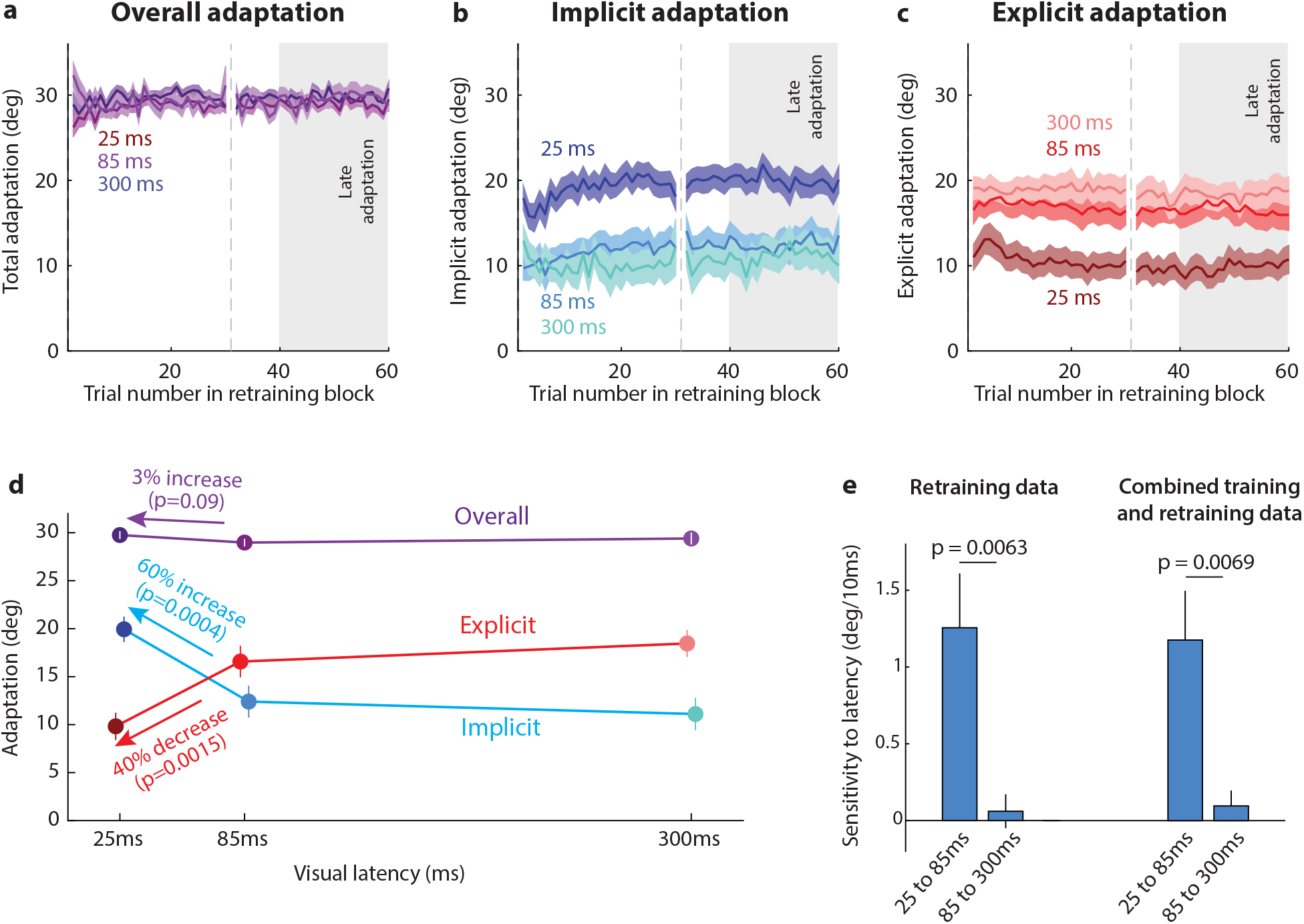
Relearning data. **(a-d)** Same as Fig. 2a-d but for the 60-trial relearning period. **(e)** Sensitivity of late learning to changes in latency. The left pair of bars compares sensitivities over the 25-85ms interval vs 85-300ms intervals for the relearning data. The right pair shows this comparison for the combined learning and relearning data. In both cases, the sub-100ms interval displays markedly higher sensitivity of implicit learning to changes in latency than the longer latency interval.

### The sensitivity of implicit sensorimotor adaptation to latency markedly increases at low visual feedback latencies

These findings reveal that short, sub-100ms visual feedback latencies can not only markedly reduce implicit adaptation, but also have a disproportionately larger effect than the higher latencies previously studied. In particular, training period data show that the 60ms decrease in latency from 85ms to 25ms results in roughly double the improvement in implicit sensorimotor adaptation than the 3-fold greater 215ms decrease in latency from 300ms to 85ms (6.6±1.9° vs 2.9±2.1°). This results in a sensitivity of adaptation per unit latency reduction that is nominally 7-fold greater for the 25-85ms interval compared to the 85-300ms interval (1.09±0.32° vs 0.14±0.10° of adaptation per 10ms of added latency, respectively (Fig. 2e)). Data from the 60-trial retraining block that followed the first generalization block, mirror the training block findings. In this case, they show that the 60ms decrease in latency from 85ms to 25ms results in triple the improvement in implicit sensorimotor adaptation than the 215ms decrease in latency from 300ms to 85ms (7.5±2.1° vs 1.3±2.4°). This results in a sensitivity that is nominally 20-fold greater over the 25-85ms interval compared to the 85-300ms interval (1.26±0.35° vs 0.06±0.11° of adaptation per 10ms of added latency, respectively (Fig. 3e)). Combining these data to obtain the most accurate comparison of the sensitivity between adaptation and latency, reveals a sensitivity of 1.18±0.32°/10ms on the 25-85ms interval compared to 0.10±0.10°/10ms on the 85-300ms interval (F(1,98)=7.6, p=0.0069, Fig. 3e). The marked contrast between these intervals underscores the exceptional sensitivity of implicit adaptation to short visual feedback latencies. Our results demonstrate that, as latency is reduced, this sensitivity increases in a rapidly accelerating non-linear trajectory. If this trajectory were to be maintained for latency reductions beyond 25ms, substantial further improvement in implicit learning might be attained for latency improvements exceeding what we were able to achieve (see Methods).

### Visual feedback latencies affect how both implicit and explicit adaptation generalize to different movement directions

We proceeded to investigate how visual feedback latencies might affect the generalization of adaptation to different movement directions, as this may provide evidence about its underlying neural representation^43–46^. During Generalization blocks performed after both the training and retraining periods, participants reached to targets in 19 different movement directions in the absence of visual feedback (-135° to 135° relative to training, spaced 15° apart in a pseudorandom order, Fig. 4a).

**Fig. 4:**
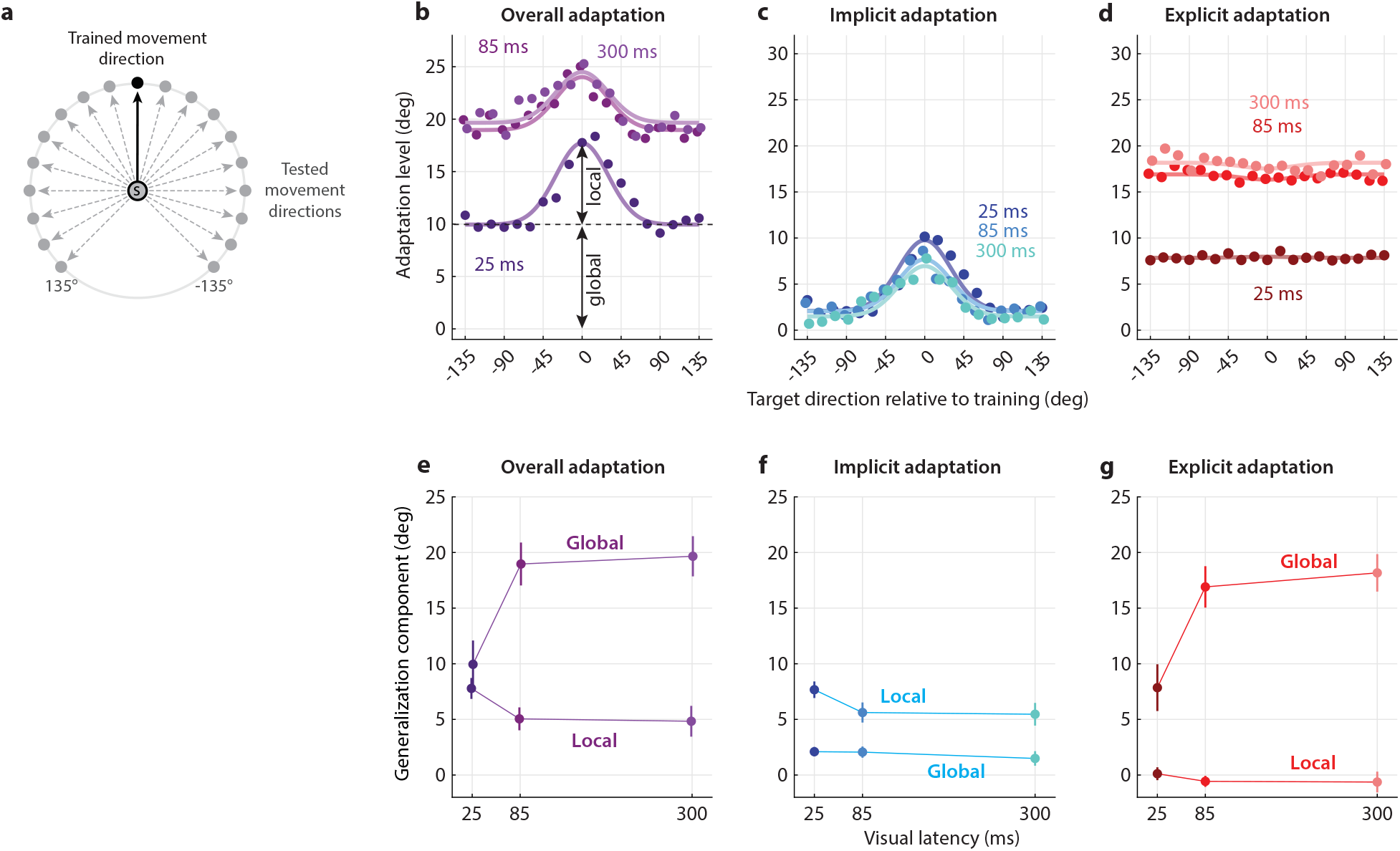
Latency reductions improve locally-generalizing implicit learning and decrease globally-generalizing explicit learning. **(a)** Generalization of VMR learning was measured with no-feedback movements across 19 different test directions that were centered on the trained target direction in 15° steps. **(b-d)** Effects of different latencies on the shape of **(b)** overall, **(c)** implicit, and **(d)** explicit generalization its locally-generalizing and globally-generalizing components. Implicit learning primarily generalizes locally and is stronger at lower latencies, whereas explicit learning primarily generalizes globally and is stronger at high latencies. **(e-g)** -generalizing and lower globally-generalizing overall learning, driven by increases in the primarily locally-generalizing implicit learning and decreases in the primarily globally-generalizing explicit learning. Error bars indicate ±SEM.

The overall across-direction generalization patterns that we observed (shades of purple in Fig. 4b) have shapes that resemble the combination of a hump-that correspond, respectively, to locally-generalizing and globally-generalizing contributions^46,47^. We thus characterized the across-direction generalization patterns as the sum of a gaussian-shaped -shaped global ; see Equation 1), by regressing each generalization pattern onto these two components. The two-associated with these regressions explain 89%, 83%, and 71% of the variance in the participant-averaged data for the overall generalization pattern for the 25ms, 85ms, and 300ms latencies, suggesting that they capture the shape of these patterns well.

When we dissected the overall generalization pattern (Fig. 4b) into separate patterns for implicit and explicit generalization (Fig. 4c-d), we found that whereas the overall pattern was comprised of sizeable contributions from both local and global components (Fig. 4e), implicit generalization was dominated by its local component and explicit generalization by its global component (Fig. 4f-g). Implicit generalization displayed local components of 5.5-8° vs global components of 1.4-2.1° for the 25ms, 85ms, and 300ms latencies, in stark contrast to local components of < 1° vs global components of 7.5-18.5° for explicit generalization. Moreover, we found that, like implicit adaptation during the training and retraining periods, the local component of implicit generalization grew as latency decreased (5.6±0.9° vs 7.7±0.7° for 85ms vs. 25ms, t(66)=1.8, p= 0.042; and 5.5±1.0° vs 7.7±0.7° for 300ms vs. 25ms, t(66)=1.7, p= 0.043). And like explicit adaptation during the training and retraining periods, the global component of explicit generalization contracted as latency decreased (16.9±1.9° vs. 7.9±2.1° for 85ms vs. 25ms, t(30)=3.2, p=0.0015; and 18.2±1.7° vs. 7.9±2.1° for 300ms vs. 25ms, t(30)=3.8, p=0.00030).

Together, our findings demonstrate that even short sub-100ms latencies can dramatically alter the internal composition of sensorimotor learning by decreasing implicit adaptation and increasing explicit strategy. Moreover, we find that implicit sensorimotor learning is far more sensitive to sub-100ms latencies than larger latencies, indicating that latency reduction within the sub-100ms range can substantially improve implicit learning.

## Discussion

Here we investigated the effects of short, sub-100ms visual feedback latencies on implicit and explicit sensorimotor adaptation. Specifically, we compared the learning and generalization of a visuomotor rotation when visual feedback was presented with a short, optimized 25ms latency vs an intermediate sub-100ms latency (85ms) and a longer 300ms latency that provided a reference condition comparable to the long latencies examined in previous studies that dissected implicit and explicit sensorimotor learning. We found that, during learning, reducing latency by just 60ms, from 85 to 25ms, led to a dramatic 50% increase in implicit adaptation and a complementary 40% decrease in explicit adaptation. Remarkably, the sensitivity of implicit adaptation to changes in latency was several-fold greater for short latencies (reducing latency from 85ms to 25ms) compared to longer latencies (reducing latency from 300ms to 85ms). This highlights the outsized importance of sub-100ms latencies in driving increased implicit sensorimotor adaptation relative to more commonly studied longer latencies. These effects were remarkably consistent when the effects of extended training were examined in the subsequent relearning period, whereby reducing latency from 85ms to 25ms led to a 60% increase in implicit adaptation, with the corresponding sensitivity to latency again several-fold greater for short vs. long latencies. Secondarily, we examined the directional generalization of learned adaptation under each latency condition, and found implicit sensorimotor learning to be dominated by locally-generalizing learning that is sensitive to latency, with explicit learning instead dominated by globally-generalizing learning. This indicates that our dissection of overall adaptation identifies implicit and explicit components with very different underlying representations. Together, our findings reveal the exquisite sensitivity of sensorimotor adaptation to visual feedback latencies in the sub-100ms range, and suggest that previously reported across-study differences in implicit learning may often be due to differences in visual feedback latency during training, underlining the need to widely measure, report, and ultimately minimize these latencies.

### Previous work on the effect of feedback latencies on motor adaptation

Most studies examining the effect of visual feedback latency on sensorimotor learning have studied large latencies above 200ms^13–15,21,24–31^ and often beyond 1000ms^15,27–31^. In fact, we were able to identify only four studies that examined the effect of latency on learning that used delays of 100ms or less and thus could have potentially examined sub-100ms latencies (and we could find no additional studies with delays below 200ms). Three of these four, however, neglected to measure the any of the actual latencies in their experimental conditions, and so the delayed condition, the baseline condition, or both could have had latencies above 100ms, rather than below it^20,22,23^. The fourth study^19^ measured a latency of 36ms at baseline, so that the 60ms delay they studied corresponded to a 96ms latency. However, despite a close correspondence to the 25ms and 85ms latencies studied here, they found no effect of delay. But because they did not dissect adaptation into implicit and explicit components, the only available measure was overall learning where we also find no effect of delay. Consequently, the details of this result are in line with what we find, despite a robust effect in our data when implicit and explicit adaptation are dissected out, because the 60ms delay we levied had opposing effects on implicit and explicit contributions that largely cancelled out when overall learning was measured. It should be noted, however, that this study attempted to isolate implicit learning by requiring short 200ms reaction times, which can reduce explicit strategy^48–51^, but the effect of this manipulation may have been tempered by the endpoint-only task feedback employed, which has the opposing effect of increasing explicit strategy^41,52,53^.

For the three studies that examined sub-100ms delays but did not measure actual latencies, it would be useful if we could make accurate post-hoc estimates of the baseline latencies. The first of these studies^21^ used the analog signals from x and y potentiometer-based sensors fed directly into an oscilloscope for visual feedback. Because oscilloscopes from the era of this study had CRT displays, the latency would likely be limited by phosphor persistence and thus be rather small, likely below 10ms, although without the make and model number for the oscilloscope used or direct information about its phosphor screen coating, we cannot be certain. Potentiometer-based position sensors should have little delay unless the output signal was intentionally smoothed by adding capacitance, which was not indicated, suggesting that the baseline visual display latency could have been rather small, perhaps as low as 10ms.

The other two studies^22,23^ were more likely to have been plagued by higher baseline latencies, although we cannot be certain. In both of these studies, a signal from a screen-mounted touch sensor was fed into a computer to detect movement endpoint so that LCD shutter glasses could be triggered to open and allow for direct endpoint feedback of the hand touching the screen where the target was displayed. The shutter glasses were reported to be able to open in just 1ms, suggesting they added little to the baseline latency. However, information was provided neither about the latency of the touch sensor nor the time needed to read-in the sensor output from it nor to process the data to determine movement completion. Contemporary devices that rely on touch sensors, such as smartphones and tablets, often suffer from touch-to-display latencies of up to 100-200ms^54–56^, suggesting that the 3-decade-old the Kitazawa et al. setup – with the first study published in 1995, when touch sensors were a newer technology – may have suffered from latencies on the high end of that range or higher. Likewise, it’s difficult to be certain about the latency associated with reading-in the sensor data to the in-the-loop computer, processing it, and sending out the shutter glass command signal, but even in our highly-optimized setup, that used a modern, much faster PC, the computer in-the-loop latency likely doubled the sensor latency (see Methods).

These three studies where latency was not measured yielded conflicting results. Held et al., 1991^20^, like Tanaka et al., 2011^19^ which measured latency, found no effect of latency for imposed delays up to 60ms. However, as with the Tanaka study^19^ the Held et al. study did not dissect learning into implicit and explicit components, and so it may also have been that a latency-driven decrease in implicit learning could have been offset by an increase in explicit learning, especially as no specific efforts to abate an explicit contribution were reported. At odds with these negative results, two studies by Kitazawa et al.^22,23^ reported significant effects for 50ms delays. One of these, however, was based on data from a single monkey^23^. The other was based on a more reasonable sample size of 21 in humans^22^; however, like the Tanaka et al. and Held et al., studies, implicit and explicit learning were not dissected out of the measured adaptation. It is thus unclear to us why the results would be positive in this case, especially as our post-hoc analysis of the probable baseline latency suggests that it was likely higher than that in both the Tanaka and Held et al. studies, and this increased baseline latency should blunt the sensitivity of the adaptive response to visual feedback delays. In sum, previous work employing short 50-60ms added delays provides conflicting results about the effect of sub-100ms latencies on sensorimotor learning, likely due to a combination of unmeasured latencies for the experimental conditions and varying contributions from implicit and explicit learning that were not dissected out.

Notably, two previous studies dissected out the effects of latency on implicit and explicit sensorimotor learning^13,15^. These studies, however, examined only much larger latencies, above 200ms in both cases, and thus did not provide information on the sub-100ms latencies prevalent in human-computer interactions including experiments for studying sensorimotor learning when latency is not isolated as a variable of interest. The Brudner et al. study^15^ measured a baseline latency of 70ms, and examined a delay of 1000ms corresponding to latencies of 70 and of 1070ms for the two conditions they compared. Although they found highly significant evidence for decreased implicit learning in the higher-latency condition, the baseline condition was already dominated by explicit learning, as 70-75% of overall adaptation was explicit compared to only 25-30% for implicit. Thus their 70ms baseline condition displayed far less implicit learning than we observed in our 25ms baseline condition (25-30% vs 65-70% of overall adaptation) and, surprisingly, also displayed less implicit learning than our 85ms latency condition, corresponding to the 60ms delay in our study. The surprisingly low level of implicit learning observed in the Brudner et al. baseline condition might be explained by two factors in their experimental design known to reduce implicit and promote explicit contributions to sensorimotor learning: the use of large amplitude perturbations^57^ and endpoint-only feedback^41,52,53,58^. They also studied an even longer 5000ms delay with similar results. The Schween et al. study^13^ was similar to Brudner et al. but examined delays of 200 and 1500ms rather than 1000 and 1500ms. They reported a baseline latency of 27ms; however, this estimate was not based on direct comparison of visual feedback and live hand motion as in the current study, Brudner et al.^15^ or Tanaka et al.^15,19^. Instead, they measured the partial latency from a statement in their code calling for the sensor reading associated with movement termination and the reading of a photodiode signal that confirmed the display of endpoint feedback. This measurement is useful but excludes the latency between actual arm motion and the time at which sensor feedback of this motion is made ready to be read-in by the computer, i.e. the sensor input latency, which constitutes nearly half of the 25ms baseline latency in our setup. Like Brudner et al., the results from Schween et al., showed significant reductions in implicit learning for the high latencies they studied, with corresponding increases in explicit learning. At their baseline latency, they found a level of implicit learning (40-45% of the total learning) that was already dominated by explicit learning and far smaller than the 65-70% level we observed at 25ms. This lower than expected level is consistent with the combination of a baseline latency that was likely higher than the reported 27ms value and the use of an endpoint-only rather than continuous feedback task. Grossly, however, both the Brudner and Schween studies are consistent with our current results in that they reported a balance between implicit and explicit learning that shifts towards being more explicit-dominated as latency increases, suggesting that reductions in latency promote implicit learning, while increases promote explicit learning. The current study adds the critical information that this balance is shifted even for small latency increases in the sub-100ms range, relevant to human-computer interactions and experiments studying sensorimotor learning, and that the sensitivity of this shift to changes in latency is, in fact, far greater in this sub-100ms range than at the longer latencies at which it had previously been identified.

A recent study tried to assess the effects of reducing latency *below* the base setup latency. For each movement, Wang et al.^17^ estimated heading direction shortly after movement onset and projected this direction out to an endpoint distance where feedback was provided. This effectively reduced the feedback latency at this endpoint by the movement time for each trial. The study consistently found that adaptation was higher for their advanced feedback condition compared to their standard base latency endpoint feedback condition, but not higher than when continuous visual feedback was provided, as in the current study. For their VMR condition, the only condition where implicit and explicit learning were measured and the one which is most comparable to the current and to previous studies, they found that implicit accounted for only about 25% of the total learning for their standard feedback condition and about 35% with advanced feedback. However, this 35% of the total learning in the advanced feedback condition is far smaller than the 65-70% observed for implicit learning for low latency condition in the current study. Even in their error-clamp condition, which employed a large 15° error signal to provide a large, continuous drive for adaptation that dwarfed the modest 0.5-2° errors that drove adaptation after the first 3-5 trials in the initial training period in the current study (see the overall learning plots in Figs. 2a and 3a), Wang et al. observed asymptotic learning that either merely matched our minimum-delay condition (20°, from their online experiments) or fell short (15°, from laboratory-controlled experiments). These observations suggest that it is the reduction of latency from positive baseline levels to near-zero values, rather than the creation of “negative” latency with advanced feedback, which is likely responsible for the enhanced learning observed by Wang et al.^17^. We do note however, that complicating comparisons with our work and other studies of latency, where latencies were precisely controlled, the effective latency reduction for the advanced feedback condition varied widely with the movement time for each trial. This resulted in a wide range even for the mean latency reductions for different individuals, with the reported values ranging between 0-300ms.

### Effects of temporal latency on neural plasticity

Evidence from neurophysiology suggests that short, sub-100ms latencies can strongly disrupt the neural plasticity that underlies learning. Spike-timing dependent plasticity in cortical synapses relies on a precise temporal coupling between synaptic input and cell firing that dictates a switch between maximally positive and maximally negative plasticity in just 10-20 ms^35–39^. In contrast, it had long been thought that plasticity in the cerebellum did not require such precise temporal coincidence. Plasticity in cerebellar Purkinje cells – widely associated with error-driven sensorimotor learning^59– 62^ – is governed by paired activity of climbing fibers carrying error signals and parallel fibers carrying contextual sensorimotor information^63–65^, with the coincidence of this timing believed to be tolerant of a broad 150ms range of climbing fiber-parallel fiber latencies. This belief was based on population data from the cerebellar vermis^66,67^. However, more recent work that examined neural responses at different latencies within individual vermal neurons found that, instead, that each exhibits much tighter tuning - up to 20-30ms – around its preferred latency^40^. The previously reported broad tuning reported previously would thus be due to different cells in the population displaying different preferred latencies ranging from 0-150ms around which the tight 20-30ms plasticity window operates. Together, the current evidence suggests that the neural plasticity underlying learning, in both cortex and cerebellum, requires tight temporal coupling of neural activity that could be disrupted by latencies as small as 50-100ms, in line with the high sensitivity of sensorimotor learning to latency that we demonstrate here.

### Implications for sensorimotor learning studies

Our finding that even short latencies can dramatically reduce implicit adaptation may explain wide across-study differences in the amount of implicit adaptation, which persist even when studies with closely aligned experiment design are compared. For example, three studies using the same type and magnitude of visuomotor perturbation – a 15° clamped cursor error – for comparable durations of training and multi-target training schedules, reported implicit adaptation levels ranging from 12° to about 25°^32–34^. Our findings suggest that previous reports of low asymptotic implicit adaptation may be, at least in part, due to high visual feedback latencies in the experiment setup. Unfortunately, we cannot know the exact contribution of different setup latencies to the different levels of implicit adaptation reported, as these were not assessed in the first place. This highlights the need to widely measure and report these “baseline” latencies so that latency effects can be meaningfully accounted for when interpreting study findings. And the need to minimize these latencies to abate setup-specific learning impairments that can occur even for sub-100ms latencies. Our experience indicates that low-latency performance can be attained by optimizing software for low-latency performance including the use of direct rather than buffered graphics updates in combination with modern low-input-lag computer gaming displays. Taking latency into consideration becomes even more important with the emergence of sensorimotor learning studies that are conducted online rather than in the lab^17,68^. With each participant using their own device, it is challenging to measure latency and, crucially, inter-individual differences in device latency could inject substantial inter-individual variability into implicit and explicit sensorimotor learning measurements due to differences in participants’ devices rather than to their learning abilities.

## Materials and Methods

### Participants and consent statement

A total of 42 individuals took part in the study (average age: 22.0±4.3 y.o., 17 male, 2 left-handed, 5 ambidextrous based on the Edinburgh Handedness questionnaire^69^; all participants used their right hand for the reaching task and operated the aim-report knob with their left, see details below). Participants provided informed consent in line with the Declaration of Helsinki, and the study protocol was approved by the Harvard University Committee on the Use of Human Subjects (IRB board).

### Experiment setup and general task description

Participants sat on a chair and made 10-cm point-to-point reaching movements on a 200Hz digitizing tablet (Intuos 3, Wacom Co., Japan) white grasping a lightweight plastic handle that contained a tablet stylus. Vision of the hand and arm was occluded. Instead, participants could receive continuously updated visual feedback in the form of a cursor (2.6mm diameter) displayed on a screen mounted horizontally above the tablet (BenQ XL2411, refreshed at 120Hz, Fig. 1a). The experiment room was dimly lit to improve screen contrast and facilitate attention to the task. To the left of the setup, we placed a knob which could be used to indicate the aiming target before each reaching movement. Participants were encouraged to move quickly to reach and stop at the target within 350ms, in which case the experiment computer played a bell sound. The experiment was coded and executed using Matlab (Mathworks, Natick, MA) with the Psychtoolbox 3.1 extension for reading the digitizing tablet and for graphical output to display targets and to provide cursor feedback^70^.

### Measurement and sources of system latency

To measure system latency, we simultaneously recorded physical motion of the handle and cursor motion using a high-speed (240 fps) camera positioned to see both in the same field of view. An LED was attached to the handle to facilitate tracking its position through the video. To obtain a measurement, we rapidly and irregularly moved the handle back and forth in the lateral direction for 5-10sec, while keeping it in the camera’s field of view. We processed the video to extract both handle and cursor motion in 2D, and used cross-correlation to identify the latency between them. For our optimized setup, we found a latency of 24.8±0.2 (mean±SEM across four videos).

To estimate the effect of different setup components on the overall latency of our optimized system, we examined different hardware and software configurations. To assess the latency due to using the digitizing tablet as a source of position information, we replaced it with a low-latency computer mouse, which reduced latency by about 16ms, i.e. 60-70% of the 25ms overall latency. Despite the tablet being responsible for most of the overall latency, however, we did not substitute it with a mouse, as the tablet provided higher position precision and, importantly, provided absolute position measurements that would be reliable even if the pen were lifted and placed on a different location on the tablet. To assess the effect of the experiment environment, including execution time, we compared latency outside vs. within the experiment environment, using mouse input in both cases, finding that latency was merely 0.8ms lower outside Matlab, i.e. about 3% of the 25ms overall latency. Finally, our screen’s input lag was 4.4ms^71^ or about 18% of the overall latency. A notable finding during the process of optimizing our setup was the importance of how the software handled screen updates. We coded our experiment to use asynchronous screen updates, as we found that synchronous screen updates added an additional ∼17ms (60-70% more) to the latency.

### Implementation of increased latency conditions

For the 85ms and 300ms conditions, we imposed additional latencies of 60ms and 275ms respectively in addition to the base system latency. We validated these latencies using the video method outlined above, obtaining 85.1±1.2 and 299.6±0.4ms, respectively.

### Experiment schedule

The experiment consisted of a familiarization block (17 movement trials to one of 19 different target directions with a latency of 25ms and no visuomotor rotation (VMR)) followed by two (Experiment 1a) or three (Experiment 1b) supersets of experiment blocks in which sensorimotor learning was assessed. Within each superset, visual feedback, when provided, was delayed by a single latency (25ms, 85ms, or 300ms) that was switched between supersets so that each participant in Experiment 1a experienced two latencies and each in Experiment 1b experienced three. The ordering of presentation of the three latencies such that the probability of experiencing each latency in the first superset was 1/3 (as was the probability in the 2nd superset, and (in Experiment 1b) the probability in the 3rd one). Each superset began with 57 trials of baseline trials with no VMR (3 pseudorandomly-ordered reaches each to targets in 19 movement directions, spaced at 15° apart and centered at the direction that would subsequently be used for VMR training). Participants then performed a 16 additional no-VMR trials in the training direction followed by 120 training trials with a VMR of ±30°. Both experiments were counterbalanced so that half of the supersets trained a +30° counterclockwise VMR and half trained a -30° clockwise VMR, independently of the experienced latency. Additionally, half the participants began with a superset that trained a +30° VMR and half that trained a -30° VMR. To reduce the possibility that learning from one superset might carry over to the next, the orientation of the VMR and the direction of the trained target location were always switched between supersets. This target direction was chosen from {-75°, or +75°} in Experiment 1a and from {0°, -120°, or +120°} in Experiment 1b, all equally likely, with 0° indicating the 12 o’clock direction.

The 120-trial single-direction training block was followed by a block testing for directional generalization of the trained adaptation. This 114-trial generalization block consisted of six reaches without visual feedback to each of the 19 different directions in the 114-trial pre-training baseline block – arrayed -135° to 135° around the trained direction in increments of 15° (Fig. 4a). This was followed by a 60-trial retraining block, with the same VMR, target direction, and latency as in the training block, and then by a second 114-trial generalization block. Completing the superset, the second generalization block was followed by a 25-trial washout block, in which participants continued to make reaches to the same trained target location with the same visual feedback latency but with no VMR, i.e. the rotation turned off, to further reduce the chance of carryover of adaptation between supersets. Rest breaks, each at least one minute in duration, were given every 57 trials within the generalization blocks, and every 20-30 trials within the training and retraining blocks.

### Decomposition of adaptation into implicit and explicit components using aim reports

We assessed implicit and explicit adaptation using aim reports immediately before each movement^15,41,42^. Prior to movement initiation, participants indicated the aim point location that they thought would result in on-target cursor motion by placing an aiming marker on the screen. The aiming marker’s distance from the start position was held constant and its direction was controlled using a rotating knob by the participant’s left hand. Participants “locked” their aiming selection by clicking on the knob before each movement. Once locked, the aiming marker’s location was fixed, and it remained visible until the end of the movement.

### Data analysis

#### Measurement of reach and aim directions and dissection of adaptation into implicit and explicit components

Position data were recorded at 200Hz and differentiated to estimate movement velocity. To estimate movement onset, we first found the moment of peak movement speed, and then went backwards until we found the moment velocity first exceeded a threshold of 6.35 cm/s. Movements were fast and so the time from this movement onset point to the peak speed point averaged only 179±5ms. We defined reach direction as the direction of a vector from the participant’s position at the peak speed point, the participant’s position at the movement onset point, and then defined the relative reach direction as the difference between the reach direction and the target direction. We defined aim direction as the angle between the participant’s selected aiming point, the start position, and the target.

For training and retraining, we defined explicit adaptation as the aim direction relative to the target, and implicit adaptation as the difference between the reach direction and the aim direction as illustrated in Fig. 1c^41,42,57,72^. For generalization, we used two different methods to dissect generalization patterns into their implicit and explicit components. In the first subtype (all generalization blocks in Experiment 1a and half the generalization blocks in Experiment 1b), we isolated implicit generalization by instructing participants to aim their hand through each probe target. In the second subtype (half the generalization blocks in Experiment 1b), we measured both implicit and explicit generalization by asking participants to report their aim prior to each probe in the same way as during training/retraining trials. To combine data from +30° VMR (counterclockwise) and -30° (clockwise) VMR supersets, we flipped clockwise data prior to further analysis.

#### Dissection of adaptation into locally-generalizing and globally-generalizing components

We extracted locally-generalizing and globally-generalizing adaptation components by regressing the observed generalization pattern onto a local training-direction-centered gaussian and a constant offset^46,47,73,74^. In equation 1, the height of the gaussian *A*_*local*_, corresponds to the strength of the locally-generalizing component that is centered on the corresponding training direction θ_*tr*_ and the height of the constant offset *A*_*global*_ corresponds to the strength of the globally-generalizing component. Note that the width of the gaussian, σ, was fixed to 30° in line with previous findings^46^.

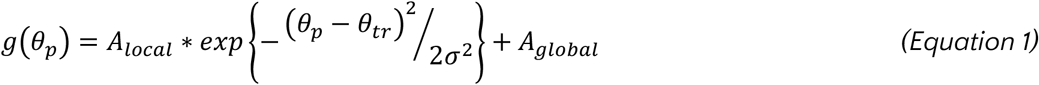

This curve was fit to each individual’s data, separately for the implicit and explicit generalization data and separately for each latency condition.

#### Baseline subtraction

To minimize the effects of any subject-specific biases and superset-to-superset carryover effects, we subtracted the average baseline from our data, separately for each individual and latency condition (i.e. within each superset), for both reaching and aiming directions (and thus for measurements of both implicit and explicit learning). For the training and retraining data, this baseline consisted of the last 10 trials before the onset of the ±30° VMR training. For the generalization data, this baseline consisted of the last pre-VMR generalization block (57 trials, three to each of the 19 tested directions).

#### Data inclusion criteria

On a small fraction of trials (0.1%), a recording issue prevented us from accurately parsing the data so that we could not determine the reach angle. We also excluded about 1% of trials as outliers with reaching angles that were more than 3 times the across-subject interquartile range away from the across-subject median of the data (separately for each latency condition). This amounted to less than 1.5% of trials (1.4±0.3%).

#### Statistical comparisons

Given that decreased implicit and increased explicit learning have been observed for larger (>200ms) latencies^13,15^, we hypothesized that the latency increases examined here would lead to decreased implicit and increased explicit learning and thus used the corresponding one-tailed t-tests to compare effects of latency.

Note that, a subset of participants (Experiment 1b) was tested under all three latency conditions, which would allow for paired comparisons; however, the remaining participants (Experiment 1a) were only tested under two out of the three latency conditions, meaning that isolating analysis to paired comparisons would exclude data. We thus used unpaired comparisons when analyzing the pooled data. We used F-tests to compare sensitivities of implicit learning to latency (Fig. 2e/3e).

## Acknowledgments

Support for this work was provided by the McKnight Scholar Award to MAS, a Sloan Research Fellowship to MAS, and a grant from NIA (R01 AG041878) to MAS.

## Competing interests statement

The authors declare no competing interests.

## Data availability

Data and code for this paper’s analyses is maintained at https://github.com/AlkisMH/Latency.

## Notes

### Competing Interest Statement

The authors have declared no competing interest.

### Summary of Updates

Minor change to title, and expansion of discussion on previous work

## Bibliography

1. Wimmer, R., Schmid, A. & Bockes, F. On the Latency of USB-Connected Input Devices. in Proceedings of the 2019 CHI Conference on Human Factors in Computing Systems 1–12 (Association for Computing Machinery, New York, NY, USA, 2019). doi:10.1145/3290605.3300650.

2. Kumcu, A. et al. Effect of video lag on laparoscopic surgery: correlation between performance and usability at low latencies. Int. J. Med. Robot. 13, e1758 (2017).

3. Anvari, M. et al. The impact of latency on surgical precision and task completion during robotic-assisted remote telepresence surgery. Comput. Aided Surg. 10, 93–99 (2005).

4. Krakauer, J. W. et al. Comparing a novel neuroanimation experience to conventional therapy for high-dose intensive upper-limb training in subacute stroke: the SMARTS2 randomized trial. Neurorehabil. Neural Repair 35, 393–405 (2021).

5. Oberhauser, M. & Dreyer, D. A virtual reality flight simulator for human factors engineering. Cogn. Technol. Work 19, 263–277 (2017).

6. Hoover, A. E. & Harris, L. R. Detecting delay in visual feedback of an action as a monitor of self recognition. Exp. Brain Res. 222, 389–397 (2012).

7. Pavlovych, A. & Stuerzlinger, W. The tradeoff between spatial jitter and latency in pointing tasks. in 187–196 (2009).

8. MacKenzie, I. S. & Ware, C. Lag as a determinant of human performance in interactive systems. in 488–493 (1993).

9. Jay, C., Glencross, M. & Hubbold, R. Modeling the effects of delayed haptic and visual feedback in a collaborative virtual environment. ACM Trans. Comput.-Hum. Interact. TOCHI 14, 8–es (2007).

10. Friston, S., Karlström, P. & Steed, A. The effects of low latency on pointing and steering tasks. IEEE Trans. Vis. Comput. Graph. 22, 1605–1615 (2015).

11. Jota, R., Ng, A., Dietz, P. & Wigdor, D. How fast is fast enough? a study of the effects of latency in direct-touch pointing tasks. in 2291–2300 (2013).

12. Brenner, E. et al. How the timing of visual feedback influences goal-directed arm movements: delays and presentation rates. Exp. Brain Res. 1–11 (2023).

13. Schween, R. & Hegele, M. Feedback delay attenuates implicit but facilitates explicit adjustments to a visuomotor rotation. Neurobiol. Learn. Mem. 140, 124–133 (2017).

14. Honda, T., Hirashima, M. & Nozaki, D. Adaptation to visual feedback delay influences visuomotor learning. PLoS One 7, e37900 (2012).

15. Brudner, S. N., Kethidi, N., Graeupner, D., Ivry, R. B. & Taylor, J. A. Delayed feedback during sensorimotor learning selectively disrupts adaptation but not strategy use. J. Neurophysiol. 115, 1499–1511 (2016).

16. de la Malla, C., López-Moliner, J. & Brenner, E. Dealing with delays does not transfer across sensorimotor tasks. J. Vis. 14, 8–8 (2014).

17. Wang, T., Avraham, G., Tsay, J. S., Thummala, T. & Ivry, R. B. Advanced feedback enhances sensorimotor adaptation. Curr. Biol. (2022).

18. Projector Central, accessed 9/27/2023. www.projectorcentral.com.

19. Tanaka, H., Homma, K. & Imamizu, H. Physical delay but not subjective delay determines learning rate in prism adaptation. Exp. Brain Res. 208, 257–268 (2011).

20. Held, R. & Durlach, N. Telepresence, time delay and adaptation. Pict. Commun. Virtual Real Environ. 232–246 (1991).

21. Held, R., Efstathiou, A. & Greene, M. Adaptation to displaced and delayed visual feedback from the hand. J. Exp. Psychol. 72, 887–891 (1966).

22. S Kitazawa, T Kohno, & T Uka. Effects of delayed visual information on the rate and amount of prism adaptation in the human. J. Neurosci. 15, 7644 (1995).

23. Kitazawa, S. & Yin, P.-B. Prism adaptation with delayed visual error signals in the monkey. Exp. Brain Res. 144, 258–261 (2002).

24. Farshchiansadegh, A., Ranganathan, R., Casadio, M. & Mussa-Ivaldi, F. A. Adaptation to visual feedback delay in a redundant motor task. J. Neurophysiol. 113, 426–433 (2015).

25. Dix, A., Helmert, J. R. & Pannasch, S. Latency in cyber-physical systems: the role of visual feedback delays on manual skill learning. in 1138–1146 (Springer, 2022).

26. Vassiliadis, P., Lete, A., Duque, J. & Derosiere, G. Reward timing matters in motor learning. Iscience 25, (2022).

27. Swinnen, S. P., Schmidt, R. A., Nicholson, D. E. & Shapiro, D. C. Information feedback for skill acquisition: Instantaneous knowledge of results degrades learning. J. Exp. Psychol. Learn. Mem. Cogn. 16, 706 (1990).

28. Dyal, J. A., Wilson, W. J. & Berry, K. K. Acquisition and extinction of a simple motor skill as a function of delay of knowledge of results. Q. J. Exp. Psychol. 17, 158–162 (1965).

29. Becker, P. W., Mussina, C. M. & Persons, R. W. Intertrial interval, delay of knowledge of results, and motor performance. Percept. Mot. Skills 17, 559–563 (1963).

30. Boulter, L. R. Evaluation of mechanisms in delay of knowledge of results. Can. J. Psychol. Can. Psychol. 18, 281 (1964).

31. Greenspoon, J. & Foreman, S. Effect of delay of knowledge of results on learning a motor task. J. Exp. Psychol. 51, 226 (1956).

32. Morehead, J. R., Taylor, J. A., Parvin, D. E. & Ivry, R. B. Characteristics of implicit sensorimotor adaptation revealed by task-irrelevant clamped feedback. J. Cogn. Neurosci. 29, 1061–1074 (2017).

33. Avraham, G., Morehead, J. R., Kim, H. E. & Ivry, R. B. Reexposure to a sensorimotor perturbation produces opposite effects on explicit and implicit learning processes. PLoS Biol. 19, e3001147 (2021).

34. Kim, H. E., Morehead, J. R., Parvin, D. E., Moazzezi, R. & Ivry, R. B. Invariant errors reveal limitations in motor correction rather than constraints on error sensitivity. Commun. Biol. 1, 19 (2018).

35. Bi, G. & Poo, M. Synaptic modifications in cultured hippocampal neurons: dependence on spike timing, synaptic strength, and postsynaptic cell type. J. Neurosci. 18, 10464–10472 (1998).

36. Froemke, R. C. & Dan, Y. Spike-timing-dependent synaptic modification induced by natural spike trains. Nature 416, 433–438 (2002).

37. Tzounopoulos, T., Kim, Y., Oertel, D. & Trussell, L. O. Cell-specific, spike timing–dependent plasticities in the dorsal cochlear nucleus. Nat. Neurosci. 7, 719–725 (2004).

38. Dan, Y. & Poo, M. Spike timing-dependent plasticity of neural circuits. Neuron 44, 23–30 (2004).

39. Zhang, L. I., Tao, H. W., Holt, C. E., Harris, W. A. & Poo, M. A critical window for cooperation and competition among developing retinotectal synapses. Nature 395, 37–44 (1998).

40. Suvrathan, A., Payne, H. L. & Raymond, J. L. Timing rules for synaptic plasticity matched to behavioral function. Neuron 92, 959–967 (2016).

41. Taylor, J. A., Krakauer, J. W. & Ivry, R. B. Explicit and implicit contributions to learning in a sensorimotor adaptation task. J. Neurosci. 34, 3023–3032 (2014).

42. Miyamoto, Y. R., Wang, S. & Smith, M. A. Implicit adaptation compensates for erratic explicit strategy in human motor learning. Nat. Neurosci. 23, 443–455 (2020).

43. Shadmehr, R. Generalization as a behavioral window to the neural mechanisms of learning internal models. Hum. Mov. Sci. 23, 543–568 (2004).

44. Gonzalez Castro, L. N., Monsen, C. B. & Smith, M. A. The Binding of Learning to Action in Motor Adaptation. PLOS Comput. Biol. 7, e1002052 (2011).

45. Rezazadeh, A. & Berniker, M. Force field generalization and the internal representation of motor learning. Plos One 14, e0225002 (2019).

46. Brayanov, J. B., Press, D. Z. & Smith, M. A. Motor memory is encoded as a gain-field combination of intrinsic and extrinsic action representations. J. Neurosci. 32, 14951–14965 (2012).

47. Zhou, W., Fitzgerald, J., Colucci-Chang, K., Murthy, K. G. & Joiner, W. M. The temporal stability of visuomotor adaptation generalization. J. Neurophysiol. 118, 2435–2447 (2017).

48. Haith, A. M., Huberdeau, D. M. & Krakauer, J. W. The influence of movement preparation time on the expression of visuomotor learning and savings. J. Neurosci. 35, 5109–5117 (2015).

49. Fernandez-Ruiz, J., Wong, W., Armstrong, I. T. & Flanagan, J. R. Relation between reaction time and reach errors during visuomotor adaptation. Behav. Brain Res. 219, 8–14 (2011).

50. Albert, S. T. et al. Competition between parallel sensorimotor learning systems. Elife 11, e65361 (2022).

51. Leow, L.-A., Gunn, R., Marinovic, W. & Carroll, T. J. Estimating the implicit component of visuomotor rotation learning by constraining movement preparation time. J. Neurophysiol. 118, 666–676 (2017).

52. Schween, R., Taube, W., Gollhofer, A. & Leukel, C. Online and post-trial feedback differentially affect implicit adaptation to a visuomotor rotation. Exp. Brain Res. 232, 3007–3013 (2014).

53. Hinder, M. R., Tresilian, J. R., Riek, S. & Carson, R. G. The contribution of visual feedback to visuomotor adaptation: how much and when? Brain Res. 1197, 123–134 (2008).

54. Kaaresoja, T. & Brewster, S. Feedback is… late: measuring multimodal delays in mobile device touchscreen interaction. in 1–8 (2010).

55. Deber, J. et al. Hammer time! A low-cost, high precision, high accuracy tool to measure the latency of touchscreen devices. in 2857–2868 (2016).

56. Ng, A., Lepinski, J., Wigdor, D., Sanders, S. & Dietz, P. Designing for low-latency direct-touch input. in 453–464 (2012).

57. Bond, K. M. & Taylor, J. A. Flexible explicit but rigid implicit learning in a visuomotor adaptation task. J. Neurophysiol. 113, 3836–3849 (2015).

58. Barkley, V., Salomonczyk, D., Cressman, E. K. & Henriques, D. Y. Reach adaptation and proprioceptive recalibration following terminal visual feedback of the hand. Front. Hum. Neurosci. 8, 705 (2014).

59. Ohmae, S. & Medina, J. F. Climbing fibers encode a temporal-difference prediction error during cerebellar learning in mice. Nat. Neurosci. 18, 1798–1803 (2015).

60. Najafi, F. & Medina, J. F. Beyond “all-or-nothing” climbing fibers: graded representation of teaching signals in Purkinje cells. Front. Neural Circuits 7, 115 (2013).

61. Tseng, Y., Diedrichsen, J., Krakauer, J. W., Shadmehr, R. & Bastian, A. J. Sensory prediction errors drive cerebellum-dependent adaptation of reaching. J. Neurophysiol. 98, 54–62 (2007).

62. Herzfeld, D. J., Kojima, Y., Soetedjo, R. & Shadmehr, R. Encoding of error and learning to correct that error by the Purkinje cells of the cerebellum. Nat. Neurosci. 21, 736 (2018).

63. Marr, D. & Thach, W. T. A Theory of Cerebellar Cortex. in From the Retina to the Neocortex 11–50 (Birkhäuser Boston, 1991). doi:10.1007/978-1-4684-6775-8_3.

64. Albus, J. S. A theory of cerebellar function. Math. Biosci. 10, 25–61 (1971).

65. Ito, M. & Kano, M. Long-lasting depression of parallel fiber-Purkinje cell transmission induced by conjunctive stimulation of parallel fibers and climbing fibers in the cerebellar cortex. Neurosci. Lett. 33, 253–258 (1982).

66. Ekerot, C.-F. & Kano, M. Stimulation parameters influencing climbing fibre induced long-term depression of parallel fibre synapses. Neurosci. Res. 6, 264–268 (1989).

67. Karachot, L., Kado, R. T. & Ito, M. Stimulus parameters for induction of long-term depression in in vitro rat Purkinje cells. Neurosci. Res. 21, 161–168 (1994).

68. Tsay, J. S., Lee, A., Ivry, R. B. & Avraham, G. Moving outside the lab: The viability of conducting sensorimotor learning studies online. ArXiv Prepr. ArXiv210713408 (2021).

69. Oldfield, R. C. The assessment and analysis of handedness: the Edinburgh inventory. Neuropsychologia 9, 97–113 (1971).

70. Brainard, D. H. & Vision, S. The psychophysics toolbox. Spat. Vis. 10, 433–436 (1997).

71. RTINGS.com, accessed 3/4/2024. https://www.rtings.com/monitor/reviews/benq/zowie-xl2411p.

72. Taylor, J. A. & Ivry, R. B. Flexible cognitive strategies during motor learning. PLoS Comput. Biol. 7, e1001096 (2011).

73. Alhussein, L. & Smith, M. A. Motor planning under uncertainty. eLife 10, e67019 (2021).

74. McDougle, S. D., Bond, K. M. & Taylor, J. A. Implications of plan-based generalization in sensorimotor adaptation. J. Neurophysiol. 118, 383–393 (2017).

